# Single Pixel Reconstruction Imaging: taking confocal imaging to the extreme

**DOI:** 10.1101/2022.11.08.515455

**Authors:** Simona Streckaitė, Dmitrij Frolov, Jevgenij Chmeliov, Andrius Gelzinis, Cristian Ilioaia, Sylvie Rimsky, Rienk van Grondelle, Leonas Valkunas, Andrew Gall, Bruno Robert

## Abstract

Light nanoscopy is attracting widespread interest for the visualization of fluorescent structures at the nanometer scale, especially in cellular biology. To achieve nanoscale resolution, one has to surpass the diffraction limit—a fundamental phenomenon determining the spot size of focused light. Recently, a variety of methods have overcome this limit, yet in practice they are often constrained by the requirement of special fluorophores, nontrivial data processing, or high price and complex implementation. For this reason, confocal fluorescence microscopy that yields relatively low resolution is still the dominant method in biomedical sciences. It was shown that image scanning microscopy (ISM) with an array detector instead of a point detector could improve the resolution of confocal microscopy. Here we review the principles of the confocal microscopy and present a simple method based on ISM with a different image reconstruction approach, which can be easily implemented in any camera-based laser-scanning set-up to experimentally obtain the theoretical resolution limit of the confocal microscopy. Our method, Single Pixel Reconstruction Imaging (SPiRI) enables high-resolution 3D imaging utilizing image formation only from a single pixel of each of the recorded frames. We achieve experimental axial resolution of 330 nm, which was not shown before by basic confocal or ISM-based systems. Contrary to the majority of techniques, SPiRI method exhibits a low lateral-to-axial FWHM aspect ratio, which means a considerable improvement in 3D fluorescence imaging of cellular structures. As a demonstration of SPiRI application in biomedical sciences, we present a 3D structure of bacterial chromosome with excellent precision.

## I. INTRODUCTION

Super-resolution fluorescence microscopy, or nanoscopy, the discovery of which was awarded the Nobel Prize in Chemistry in 2014^1^, has already led to remarkable progress in cell biology^2–4^. It is attracting increasing attention for nanometer-scale visualization of fluorescent structures, whereas theemerging improvements in this field encourage nanoscopy to generally spread in life science research as well as in other fields like synthetic (bio)material science^5,6^.

From a general point of view, achieving nanoscale resolution requires one to overcome the diffraction limit—a fundamental physical limitation, discovered by Abbe almost 150 years ago, which states that light cannot be focused by any optical system to a spot smaller than approximately half its wavelength^7^. Consequently, in conventional light microscopy, the lateral and axial values of full widths at half maximum (FWHM) of the point-spread function (PSF), which define the maximum resolution of the approach, are in the 200–400 nm and 500–800 nm ranges, respectively. These limits were circumvented by making use of photo-physical effects, computational methods, or a combination of both^8,9^. The earliest (and most successful in terms of resolution) approaches rely either on single-molecule localization microscopy (SMLM)^10,11^ or stimulated emission by depletion (STED)^12^ and exhibit a resolution of a few tens of nanometers. Unfortunately, these methods have intrinsic experimental constraints, which sometimes limit their applicability and/or decrease their resolving power. In the last decade, a series of approaches were designed, yielding lower resolution but easier to implement. Structured illumination microscopy (SIM), in which the sample is excited in a wide field through sub-diffraction grid-like structures to modulate the emission of the molecules, yields lateral resolution of about 100–130 nm^13^. Several numerical treatments of the microscopy signals have also proven successful in improving the resolution. Some examples are super-resolution radial fluctuations (SRRF) yielding *ca*. 70–150 nm lateral resolution, but being rather limited for volumetric imaging^14,15^, or super-resolution optical fluctuation imaging (SOFI), which is based on higher-order statistical analysis of temporal fluctuations and offers 5-fold resolution enhancement in 2D^16^, which makes it suitable for imaging of fluorescent blinking molecules.

For routine measurements in a cellular biology laboratory, however, many aspects of the imaging system are important, besides its intrinsic resolution. These aspects include the possibility to image living systems under relevant physiological conditions in three dimensions and to use excitation light of low enough intensity to avoid sample photobleaching. In addition, the simplicity of sample preparation, of imaging procedure and of data analysis must be taken into account. Existing nanoscopy approaches are now often implemented in large imaging facilities, and are still seldom present in individual cellular biology laboratories. To become routine, ideal super-resolution imaging methodologies should be simple, inexpensive to implement, and compatible with the conventional epifluorescence and confocal microscopes, as the latter are still currently the most popular techniques for fluorescence imaging.

In laser-scanning confocal microscopy, the diffraction-limited resolution can theoretically be improved by *ca*. 1.4 times using an infinitely small pinhole^17^. In practice, however, the pinhole size is maintained large enough to avoid signal losses and diffraction phenomena. Confocal microscopy with commonly used pinholes of several tens of micrometers displays marginal resolution increase, and its main interest resides in the sectioning ability and contrast enhancement^8,14,18–20^. Increasing the resolution of confocal microscopy was proposed a few decades ago^21^ by making use of a detector array to collect emission information. This approach, image scanning microscopy (ISM), was predicted to provide a two-fold increase in resolution upon reassignment of the detector pixels^22,23^. Experimentally, ISM was shown to yield *ca*. 200–270 nm lateral resolution^24,25^, which can be improved to 150 nm for Fourier-filtered images^24^ and to 193 nm with adaptive pixel reassignment (APR-ISM)^26^. This relatively simple approach constitutes the basis of the popular Airyscan commercial systems^27^, in which each element of the detector represents a small pinhole that is several times smaller than those used in confocal microscopy.

In this work, we review the principles of the confocal scanning microscopy and present an even simpler method to experimentally achieve the theoretical limit of the resolution of confocal microscopy by improving the ISM detection procedure. This technique—Single Pixel Reconstruction Imaging (SPiRI)—can be implemented in a standard camera-based laser-scanning fluorescence imaging systems and straightforwardly applied for high-resolution 3D imaging. Its resolution is predicted to be 30% higher than that of classical confocal microscopy in all three dimensions, a value which we experimentally demonstrate by obtaining the PSF of a home-made SPiRI imaging system using small fluorescent beads. Its excellent lateral-to-axial resolution ratio, which, to our knowledge, was not experimentally shown before for traditional confocal or basic ISM imaging systems, is particularly effective for 3D imaging, and we illustrate this by providing a 3D reconstruction of an *Escherichia coli* (*E. coli*) chromosome, stained with a fluorescent DNA intercalator. This approach, which is here applied only to fluorescence imaging, can be directly transferred to any type of emission microscopy, opening the way for achieving similar increases in resolution when using these techniques.

## II. METHODS

### Microscopy

A commercial inverted epi-fluorescence microscope (Ti-U, Nikon, Champigny-sur-Marne, France) equipped with a 100x CFI PLAN APO objective lenses (Nikon, Champigny-sur-Marne, France) was coupled to an iXon Ultra 978 EMCCD camera (Andor, Belfast, UK), via a 2x Optomask (Cairns Research, Faversham, UK). Type F immersion oil (Olympus, Japan) was used for the objective lens. The excitation wavelength was provided by a 488 nm LX OBIS laser (Coherent, Les Ulis, France) and the beam collimated with a spatial filter assembly (KT310, LA1986-A; Thorlabs, Maisons Laffitte, France). Fluorescence emission was collected through the aforementioned microscope system equipped with an excitation filter (FL488-10; Thorlabs, Maisons Laffitte, France), a dichroic mirror (ZT488rdc; Chroma Technology; Vermont, USA) and an emission filter (GFP525-39; Thorlabs, Maisons Laffitte, France). Samples were mounted between two sealed coverslips and attached to a XY-translation stage (P-733.2CD; Physik Instrumente, Aix en Provence, France) by a home-built sample holder. In-house computer scripts controlled a DAC communications interface (PCU-100; Andor, Belfast, UK) to coordinate laser emission, the XYZ-nanopositioning system (E-725.3CD, P-733.2CD, P-725.4CD; Physik Instrumente, Aix en Provence, France) and camera acquisition. Typical exposition times were 15 and 7 milliseconds per point during the fluorescing bead and *E. coli* nucleoid measurements, respectively. Emission of these samples was collected within 505–545 nm window.

### Image processing

Recorded SPiRI images were analysed by custom-written software and post-processed with open-source image analysis platforms Fiji^28^ and Icy^29^. Deconvolution was carried out with a plug-in^30^ implemented in Fiji by using the Richardson-Lucy algorithm with total-variation regularization^31,32^. Each 2D plane was deconvolved by using the regularization parameter set to 10^6^, and 10 iterations were performed during the algorithm run with experimentally obtained PSF (see text). Volumetric 3D reconstructions from deconvolved 2D planes were performed with a Volume Viewer (Kai Uwe Barthel, Internationale Medieninformatik, HTW Berlin Berlin, Germany) plug-in implemented in Fiji.

### Bead measurements

Solutions of yellow-green (505/515) fluorescent carboxylated microspheres (F-8795; Fisher Scientific, Illkirch, France) of different sizes were diluted 10^4^ fold with ethanol. A 5 μl drop of the solution was spread over a coverslip and allowed to dry, yielding a low-density bead monolayer on the coverslip.

### Bacterial strain, growth conditions, nucleic acids staining procedure and sample preparation

*Escherichia coli* HupAB cells were grown in Luria Broth (LB) medium overnight at 30°C and at 160 rpm from single-colony isolates before being diluted. After 24 hours, the cultures were diluted 1:1000 and grown further until they reached the mid-exponential phase (OD_600_ = 0.4 to 0.5). 500 μl of culture was centrifuged and the pellet re-suspended in phosphate buffer (PBS). This procedure was repeated two times. Cells were then fixed for 20 min with a 3% formaldhehyde solution and subsequently washed a few times with PBS. Staining of the nucleic acids was performed by incubating the fixed cells for 20 minutes (in dark, on ice) with SYBR® Gold (Fisher Scientific, Illkirch, France) stain. Aliquots of 5 μl were deposited on 1.5% agarose pads mounted on a 22 μm circular microscope coverslip (Ref. 011620; Paul Marienfeld, Lauda-Königshofen, Germany) fixed to a 1 mm deep silicone O-ring. After a few minutes, the sample was “sandwiched” by the addition of the second coverslip.

## III. IMAGE FORMATION IN OPTICAL MICROSCOPY: RECALLING THE BASICS

The first step from conventional wide-field microscopy towards breaking the diffraction limit was made with an introduction of the scanning microscopy^33^. Precision of the currently achievable mechanical movement is on the nanometre level or even below. Thus, as long as one controls variations in space that occur, for example, due to translational movement of the sample, these changes can be recorded optically and then converted into higher-resolution images. This idea is one of the cornerstones of the SPiRI technique presented in this work. Before discussing this newly proposed method, however, let us briefly recall the basic principles of the image formation in the optical scanning microscopy.

### A. Intensity distribution of the focused excitation beam

Spatial distribution of the excitation intensity in the sample around the focus of the optics is usually described according to the Huygens–Fresnel principle, according to which the amplitude of the electric field at point 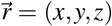 can be expressed as

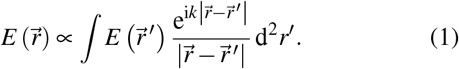

Here the integral is taken over the incoming wavefront confined by the aperture and 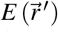 is the amplitude of the electric field of the light wave at the point 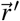 of aperture of the lens. If the sample is located in the medium with refractive index *n*, the wavevector is *k* = 2*πn*/*λ*_exc_, *λ*_exc_ being the wavelength of the laser in vacuum.

Eq. 1 is significantly simplified when the angle *ϑ*, at which the aperture is seen from the focus (see Fig. 1a), is small and assuming the plane incoming wavefront, *i. e*. 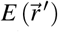 = const in the plane of the aperture. If the focal length *F* is much larger than the distance between the point 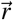 and the focus (*i. e. x, y, z ≪ F*), the integral in Eq. 1 can be calculated analytically, and when the point 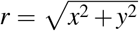 belongs to the focal plane (blue point in Fig. 1a),

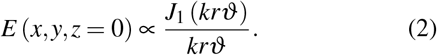

Here 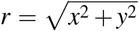 and *J*_1_ (*ξ*) is the first-order Bessel function of the first kind. This leads to well-known expression of the distribution of the focused light intensity around the focal point:

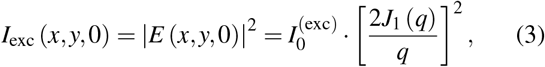

where 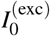 is the intensity at the focal point *x* = *y* = *z* = 0 and

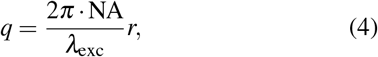

with NA = *n* sin *ϑ* denoting the numerical aperture of the focusing optics. This expression reasonably describes the intensity distribution in the focal plane even when angle *ϑ* is not very small (up to *ϑ ≈* 30°). Note that the irradiance distribution of linearly polarised light in the focal plane is no longer radially symmetric. However, even in such a case, the resulting difference between the actual FWHM of the irradiance distribution in the *x* and *y* directions and the one obtained from Eq. 3 (for which FWHM_*x,y*_ = 0.5145*λ*_exc_/NA) does not exceed 10%^34^.

**Figure 1.**
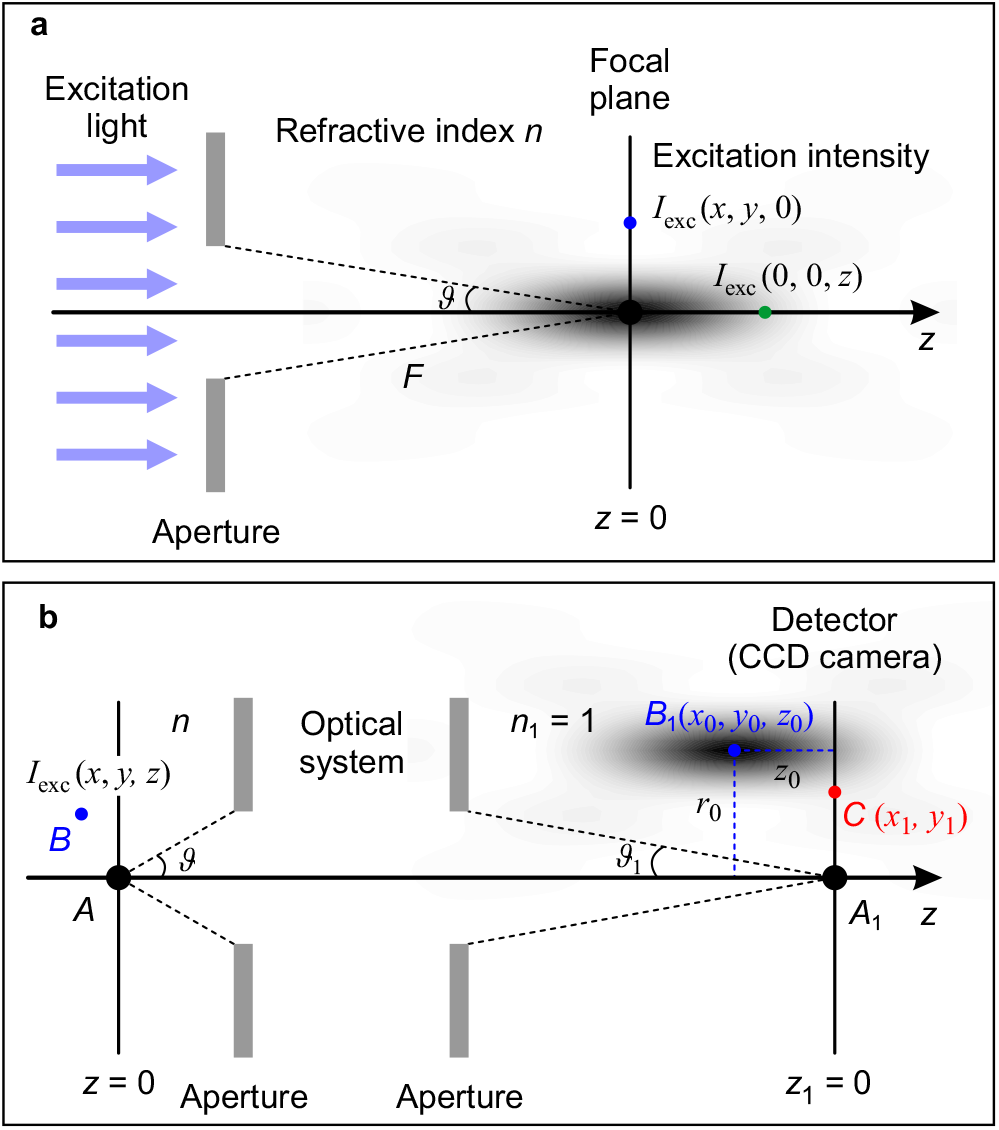
**(a)** Intensity distribution of the focused excitation beam around the focus, see text for details. **(b)** Registration of the fluorescence from a single emitter (point *B*) at the detector plane (*z*_1_ = 0), see text for details.

In the same limiting case (small angle *ϑ* and large *F*), a simple analytical expression of the irradiance distribution along the optical axis can also be obtained (green point in Fig. 1a):

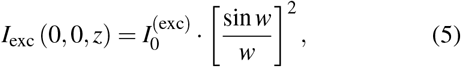

here

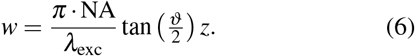

The FWHM of this distribution can be written as FWHM_*z*_ = 0.8859*λ*_exc_/(NA·tan(*ϑ* /2)), which is several times larger compared to the width of the radial distribution.

### B. Intensity distribution of the focused emission signal

In the case of a single fluorophore randomly oriented at the point 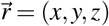, the probability for this molecule to absorb a photon is proportional to the intensity of the excitation beam at this point 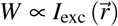. When this molecule is located precisely at the focus of this beam (point *A* in Fig. 1b), its emission is focused to the point *A*_1_ on the other side of the optical system, thus producing a radial intensity distribution at the detector plane, similar to the one described by Eq. 3:

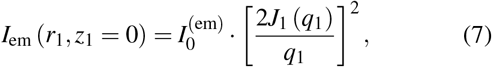

here

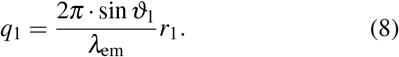

The optical design of the microscope usually obeys Abbe’s sine condition *n* sin *ϑ* = sin *ϑ*_1_·*M*, where *M* is the lateral magnification of the microscope. Thus *q*_1_ can be rewritten in terms of the numerical aperture of the objective:

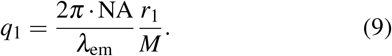

When the molecule is shifted out of focus by a small (compared to the focal length) distance and is now located at point *B* with coordinates (*x, y, z*), its fluorescence signal will produce a three-dimensional irradiance distribution 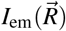 around the point *B*_1_ whose coordinates are (*x*_0_ = *xM, y*_0_ = *yM, z*_0_ = *zM*^2^/*n*) with respect to the central point *A*_1_. As a result, the detected fluorescence intensity at some point *C* (*x*_1_, *y*_1_) in the detector plane *z*_1_ = 0 will be equal to the products of *I*_em_ (*x*_1_ − *x*_0_, *y*_1_ − *y*_0_, *z*_0_) and the probability *W* for the molecule to be excited by the laser pulse, thus

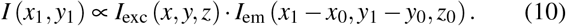

When the molecule remains in the focal plane of the laser-focusing optics (*z* = 0), this expression can be simplified:

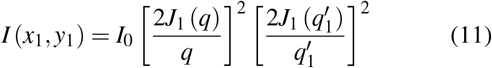

with

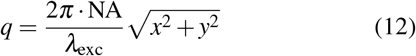

and

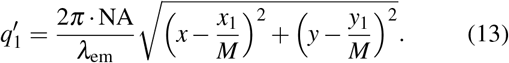

Alternatively, if the molecule remains on the optical axis, Eq. 10 simplifies to

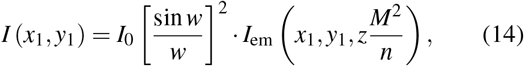

where *w* is given in Eq. 6.

### IV. SINGLE PIXEL RECONSTRUCTION IMAGING

In confocal microscopy, as discussed above, the FWHM of the radial intensity distribution *I*_exc_(*x, y*) given in Eq. 3 is

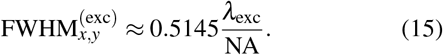

The light emitted by a single excited fluorophore and focused on the CCD camera will display a similar intensity distribution *I*_em_(*x, y*). After rescaling the obtained image while taking into account the lateral magnification of the microscope, its FWHM will be

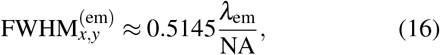

where *λ*_em_ is the dominating emission wavelength. Usually, *λ*_em_ > *λ*_exc_, therefore *I*_em_(*x, y*) is wider than *I*_exc_(*x, y*).

The amplitude of the emission intensity distribution will be proportional to the probability for the molecule to be excited by the laser, which in turn depends on the excitation light intensity at that point (*cf*. color profiles in the middle part of Fig. 2a). When a point fluorophore is moved within the focal plane, its fluorescence intensity will change according to the laser intensity at the position of the molecule, while its image will move on the detector array. If the size of the detector is small relative to the image of this point fluorophore, the amplitude of the fluorescence recorded by this detector will change more than the amplitude of the excitation beam profile. Such a small detector can be a pixel of the CCD camera, when the light intensity does not vary significantly over the entire pixel. The image of a region of interest (ROI) is constructed by using at each step the registered intensity with this small detector. When a single emitter is located at some point with coordinates (*x, y, z* = 0), let us analyze the intensity detected *just in the central pixel* of the CCD camera (point *A*_1_ in Fig. 1b and shaded pixel in the middle part of Fig. 2a). If this pixel is small enough, by substituting *x*_1_ = *y*_1_ = 0 into Eq. 13 we obtain:

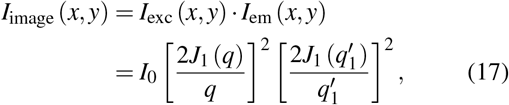

where 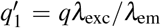 and *q* is given in Eq. 12. In other words, this method, which actually corresponds to confocal microscopy with a nano-sized detector, yields the absolute theoretical limit of the confocal microscopy (corresponding to nanometer-sized pinholes). Being the product of the two axially symmetric distributions, Eq. 17 defines a narrower distribution; its FWHM is determined by both the laser excitation wavelength and the dominating emission wavelength of the molecule:

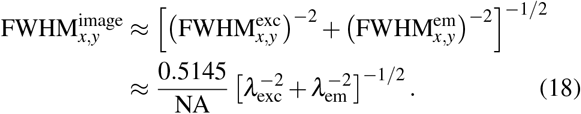

Numerical simulations of the SPiRI method applied to a single point emitter are shown in Fig. 2b. In conventional wide-field fluorescence microscopy, the registered distribution (Fig. 2b, grey shaded area) of *I*_em_(*x, y*), would be mainly determined by the emission wavelength *λ*_em_ (for *λ*_em_ = 520 nm and NA = 1.45, FWHM_*x,y*_ = 185 nm). In confocal scanning microscopy (when the image is recorded by a photomultiplier), the distribution *I*_exc_(*x, y*) will be somewhat narrower, determined by the laser excitation wavelength *λ*_exc_ (for *λ*_exc_ = 488 nm and the same NA = 1.45, FWHM_*x,y*_ = 173 nm). By contrast, SPiRI results in the discretized form of Eq. 17 that represents an even narrower distribution than either *I*_exc_ or *I*_em_. The pixel size of the obtained image is equal to the step size of the scanning procedure, and for the conditions described above, FWHM_*x,y*_ = 126 nm. As shown in Fig. 2d, such an approach can distinguish two point emitters separated by only 160 nm. The inset of Fig. 2b shows that using a larger detector leads to the situation corresponding to that of a large physical pinhole of confocal microscopy, which results in a significant drop in resolution.

**Figure 2.**
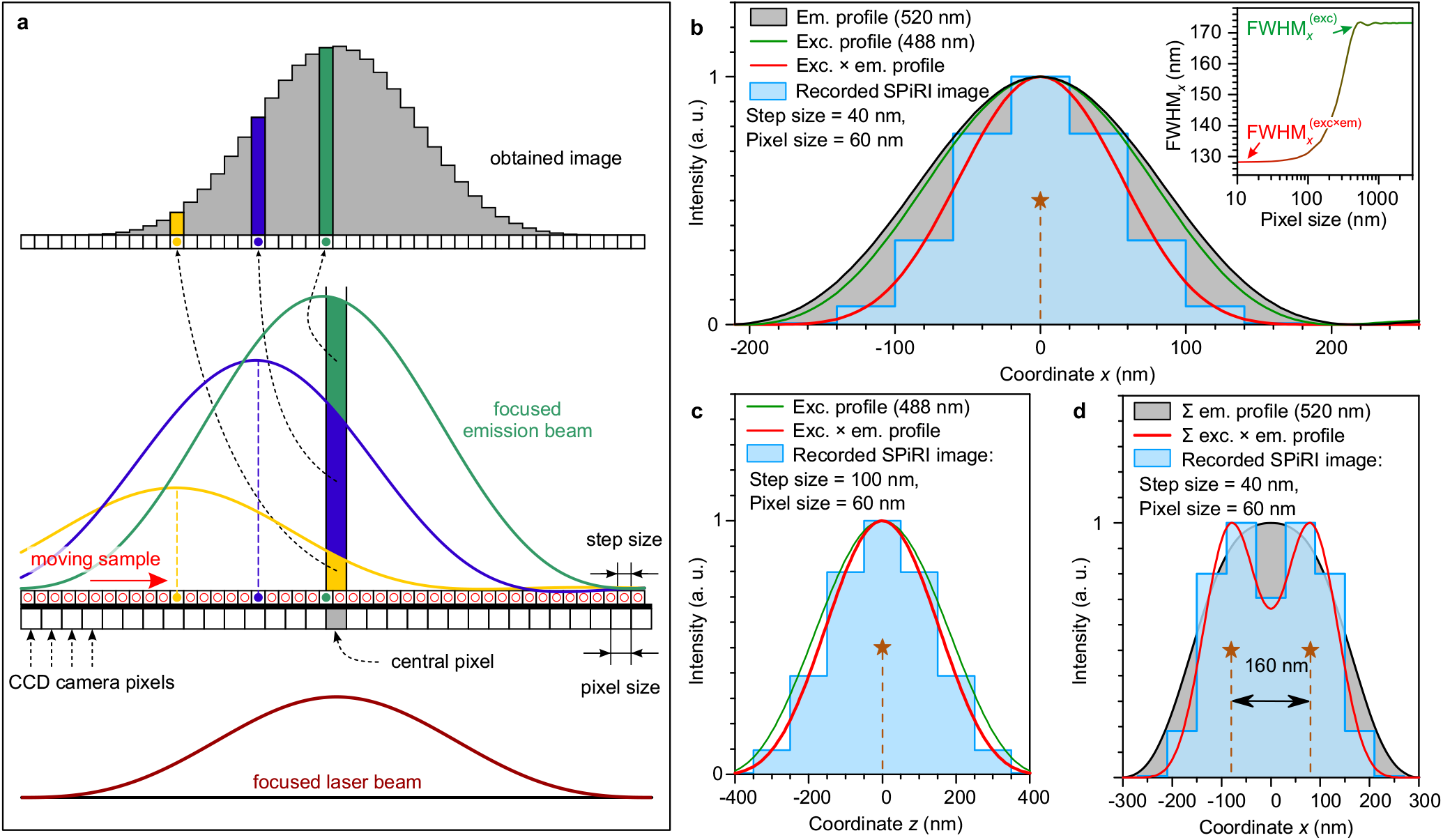
Single Pixel Reconstruction Imaging in a nutshell. **(a)** The principal scheme of the SPiRI data recording and image reconstruction for a one-dimensional (1D) case. The sample is scanned with a focused laser beam with a step size of tens of nanometres, and the final high-resolution image is reconstructed by assigning the value of the recorded intensity only from a single pixel (the one corresponding to the peak in the focused laser beam profile) of each of the recorded camera frames. The result of the scanning step size on the obtained reconstructed image is discussed in the Supporting Information (SI) (see Oversampling) and shown in Fig. S1. **(b)** Simulation of a single point emitter in 1D: emission profile in a conventional microscopy (grey area), intensity profile of the focused excitation laser beam (green line) and the reconstructed SPiRI image, obtained by using scanning step of 40 nm (blue area). SPiRI image corresponds to the discretized excitation *×* emission profile (red line). **Inset** shows the dependence of the FWHM of the simulated image on the scanning pixel size. **(c)** Simulation of the axial distribution (*z* direction) of the SPiRI-recorded image of a single point emitter, obtained by scanning along the optical axis with step size of 100 nm (blue area). **(d)** Simulation of the lateral fluorescence intensity distribution from two point emitters separated by 160 nm: emission profile after laser excitation exhibits a single flat top (grey area), while in the SPiRI-reconstructed image both emitters can be clearly distinguished, due to a significant (∼32%) intensity drop between the two observed peaks. All the simulations were performed assuming *λ*_exc_ = 488 nm, *λ*_em_ = 520 nm, and NA = 1.45, corresponding to our experimental conditions.

Along the *z* axis, the axial FWHM can be determined analogically, by moving a single emitter along the optical axis. In such a case from Eq. 14 we obtain:

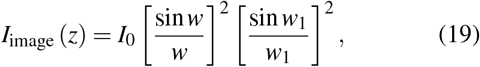

where

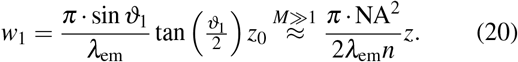

The FWHM of this distribution is

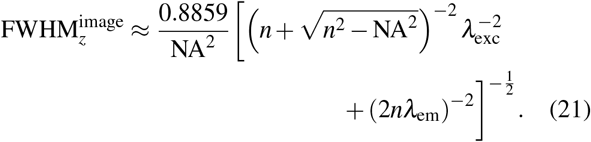

The numerical simulations for the single emitter moving along the axial direction are shown in Fig. 2c. For the same wavelengths as above and a sampling area of 60 nm *×* 60 nm on each CCD camera pixel, we obtain FWHM_z_ = 345 nm. In confocal scanning microscopy with an infinitely large pinhole one would get 397 nm, see green line in Fig. 2c.

To summarize, for *λ*_em_ = 520 nm, *λ*_exc_ = 488 nm and NA = 1.45, application of SPiRI to a single-point emitter theoretically results in FWHM_*x,y*_ = 126 nm and FWHM_z_ = 345 nm, while confocal microscopy results in significantly larger values of FWHM_*x,y*_ = 185 nm and FWHM_z_ = 398 nm.

In real measurements, the image obtained using a microscope objective is further enlarged—this allows one to choose the magnification factor of the optical system and the pixel size of the CCD detector in such a way that once projected to a CCD camera, the registered image of a point fluorophore spreads over a minimum of three pixels in each detector dimensions. Then, one of these pixels (corresponding to the peak position of the focused laser beam) of the CCD camera can be used as the above-mentioned effectively nano-sized detector required by SPiRI, although its actual size might be considerably larger.

## V. CHARACTERIZATION OF THE POINT-SPREAD FUNCTION

### A. Constructing the image of multiple emitters

The SPiRI method is not only extremely simple to implement, but it is also compatible with all image post-processing techniques utilized in confocal microscopy, such as deconvolution using the PSF^35^. Indeed, the numerical values for the FWHM of the registered image in the lateral and axial directions (shown in Fig. 2b and c) were obtained from simulating the signal of a single point emitter, and represent the theoretical limit for the resolution of the SPiRI technique. In a more realistic case involving multiple emitters, it can be assumed (using a one dimensional case for simplicity that is illustrated in Fig. 3) that the spatial excitation laser profile follows some function *P* (*x*) (centered at its maximum position), and the emission profile of a single point emitter on the CCD camera plane is given by some other function *F* (*x*) (again, centered at its maximum, i. e. the position of the emitter). The concentration of the point emitters in the sample is given by some distribution function *S* (*x*). Upon the movement of the sample across the laser excitation profile, for each position *x*_0_ of the sample, the intensity at various coordinates *x* will be registered, and the intensity distribution function *I* (*x*_0_, *x*) will depend on both these coordinates.

**Figure 3.**
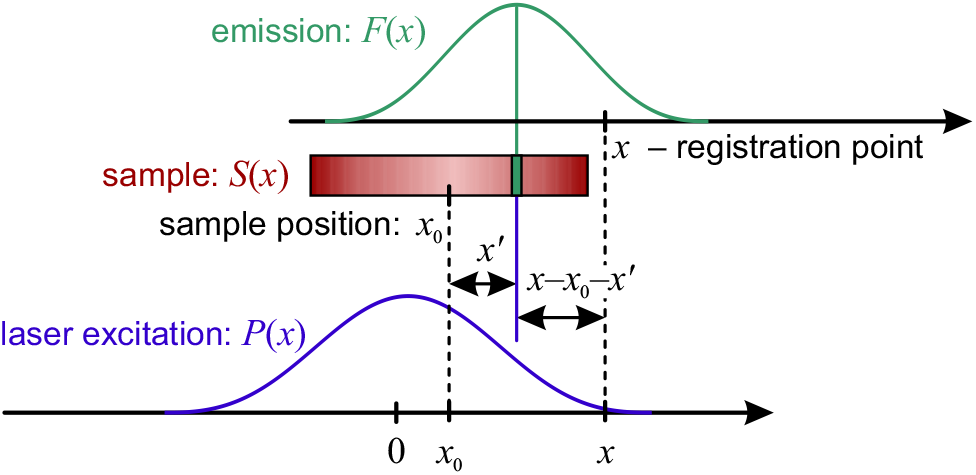
Registration of the fluorescence from the one-dimensional extended light emitter. *P* (*x*) denotes the laser excitation intensity profile in the focal plane of the objective, *F* (*x*) – the intensity distribution of a single point emitter as registered on the CCD camera, and *S* (*x*) stands for the distribution of the point emitters within the sample. See text for more details.

The emission from a point located at *x*^*′*^ (with respect to the central position of the sample) creates a signal at some point *x* of the CCD camera that is proportional to the product of the laser excitation intensity *P* (*x*_0_ + *x*^*′*^), concentration of the molecules at that point within the sample *S* (*x*^*′*^), and the emission profile *F* (*x* − *x*_0_ − *x*^*′*^). Therefore, the total intensity at *x* can be calculated by integrating over the whole sample:

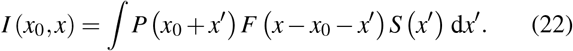

This expression, although similar to the convolution of some point-spread function (PSF) with the source function *S* (*x*^*′*^), is not exactly of the mathematical form of convolution. However, at the central pixel of the CCD camera (where *x* = 0), it yields the usual definition of convolution:

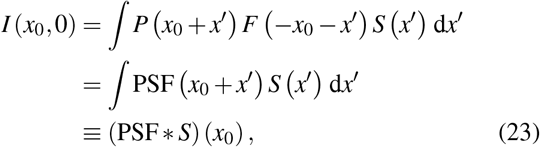

here we denoted PSF(*x*_0_) = *P* (*x*_0_) *F* (−*x*_0_). For symmetrical emission profile (i. e. if *F* (−*x*) = *F* (*x*)), we obtain the simple relation PSF(*x*_0_) = *P* (*x*_0_) *F* (*x*_0_), which is just a 1D case of the more general expression in Eq. 17.

In the ideal case, the PSF could be determined as the product of the light intensity distributions of the focused excitation and emission signals. The analytical expression of these distributions in terms of the Bessel or sinc functions can be obtained only in the simplest cases discussed above, assuming mostly co-axial light propagation, ideal focusing, no aberrations, etc. The realistic PSF must be determined experimentally by obtaining the image of a point emitter—a small enough nm-sized fluorescent object. Simulations of the images of “flat beads” of different sizes, performed for *λ*_exc_ = 488 nm and *λ*_em_ = 520 nm (see Fig. S2), revealed that all beads with diameter smaller than 50 nm produced the same SPiRI images, and can thus be considered as a single point emitters.

### B. Experimental measurements of the PSF

The experimental PSF of the SPiRI microscopy was obtained by imaging the 47-nm-sized fluorescing beads. Beads were scanned with 20-nm *x & y* steps, and 100-nm *z* steps, and the final image was reconstructed by using either the whole area detector (equivalent to confocal microscopy, see Fig. 4a) or only the central pixel (SPiRI, see Fig. 4b). A ∼30% gain in lateral and axial resolution is observed for SPiRI as compared to confocal microscopy (Fig. 4c), in agreement with the theoretical predictions. The lateral and axial cross-sections of the resulting 3D PSF (see Fig. 4e), obtained using SPiRI (Fig. S3), yield FWHM of 165, 182, and 329 nm in the *x, y*, and *z* directions, respectively (see also Table S1), showing that the method yields an extremely low axial/lateral FWHM aspect ratio, which on average is around 1.7.

**Figure 4.**
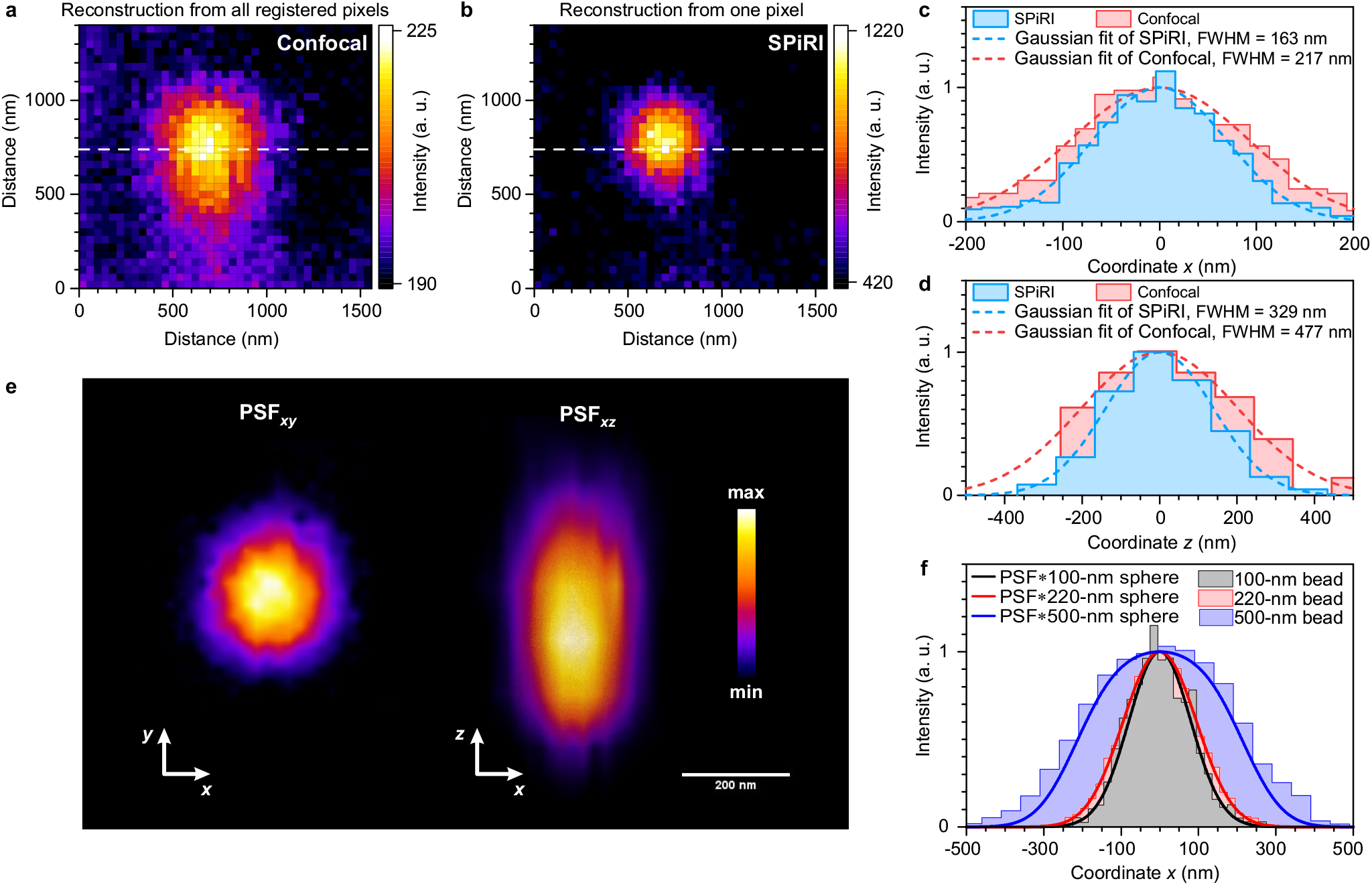
Bead measurements using SPiRI and determination of the experimental PSF. **(a)** Reconstructed image of a 47-nm bead, obtained by summing over the 30 *×* 30 pixel detector area (as in confocal microscopy). **(b)** Reconstructed image of a 47-nm bead, obtained by the SPiRI method. **(c)** Horizontal cross-sections of the reconstructed images in panels (a) and (b) along the white dashed line (red and blue shaded areas, respectively) and their respective Gaussian fits (dashed lines). **(d)** Axial distribution of the fluorescence intensity from the 47-nm bead, obtained by using SPiRI and confocal techniques (red and blue shaded areas, respectively) and their respective Gaussian fits (dashed lines). **(e)** Cross-sections in the *xy* and *xz* planes of the three-dimensional PSF of the SPiRI system, obtained from the reconstruction of the 47-nm bead images. **(f)** Lateral cross-sections of the SPiRI images of variously sized beads (shaded areas), compared to the numerical convolution of the three-dimensional Gaussian PSF (FWHM_*xyz*_ = 180 : 180 : 330 nm) from panel (e) with a three-dimensional sphere of a corresponding bead size (solid lines). Experimental cross-sections in lateral and axial directions (panels (c) and (d)) were fitted with the Gaussian functions, thus also obtaining the mean background intensity level. This background was subtracted and the resulting experimental distributions were rescaled so that their Gaussian fits were normalized to the 0–1 interval. For panels (a)–(d), a 47-nm bead was scanned by 20-nm *x & y* steps, and 100-nm *z* steps; 3D reconstruction of PSF obtained from 9 planes in the z direction. For panel (f), 100- and 200-nm beads were scanned by 20-nm *x & y* steps, and a 500-nm bead was scanned by 47-nm *x & y* steps.

Validation of the PSF was achieved by convolving the Gaussian PSF (for simplicity, using FWHM_*x*:*y*:*z*_ = 180 : 180 : 330 nm) with the spheres of 100, 220, and 500 nm in diameter. The calculated intensity distributions in the lateral cross-section exhibit a very good agreement with the actual experimental results of differently-sized beads, see Fig. 4f. An example of the deconvolution procedure using an experimentally obtained PSF, resulting in a recovery of an actual size of 100-nm bead, is shown in Fig. S4.

## VI. EXAMPLE OF 3D IMAGING BY SPiRI: *E. coli* NUCLEOID

The SPiRI approach is in principle applicable to any type of emitting sample. As an example of an application of the method, images were obtained for the *E. coli* bacterial nucleoid stained with SYBR® Gold, a cyanine-based fluorescent nucleic acid intercalator. Whole *E. coli* cells were scanned with 40-nm *x & y* steps, and 200-nm *z* steps using a 488-nm laser excitation; the SYBR® Gold emission was collected at 505–545 nm region (see Fig. 5a and Fig. S5). While confocal microscopy could barely distinguish details in this structure, SPiRI images reveal many more details (see intensity profiles in Fig. 5b and Fig. S6). To enhance these structural features, each image along the z axis was deconvolved using the experimentally determined PSF (Fig. 5a). In Fig. S6 we show that even after the deconvolution procedure, confocal images are not as detailed as raw SPiRI images.

**Figure 5.**
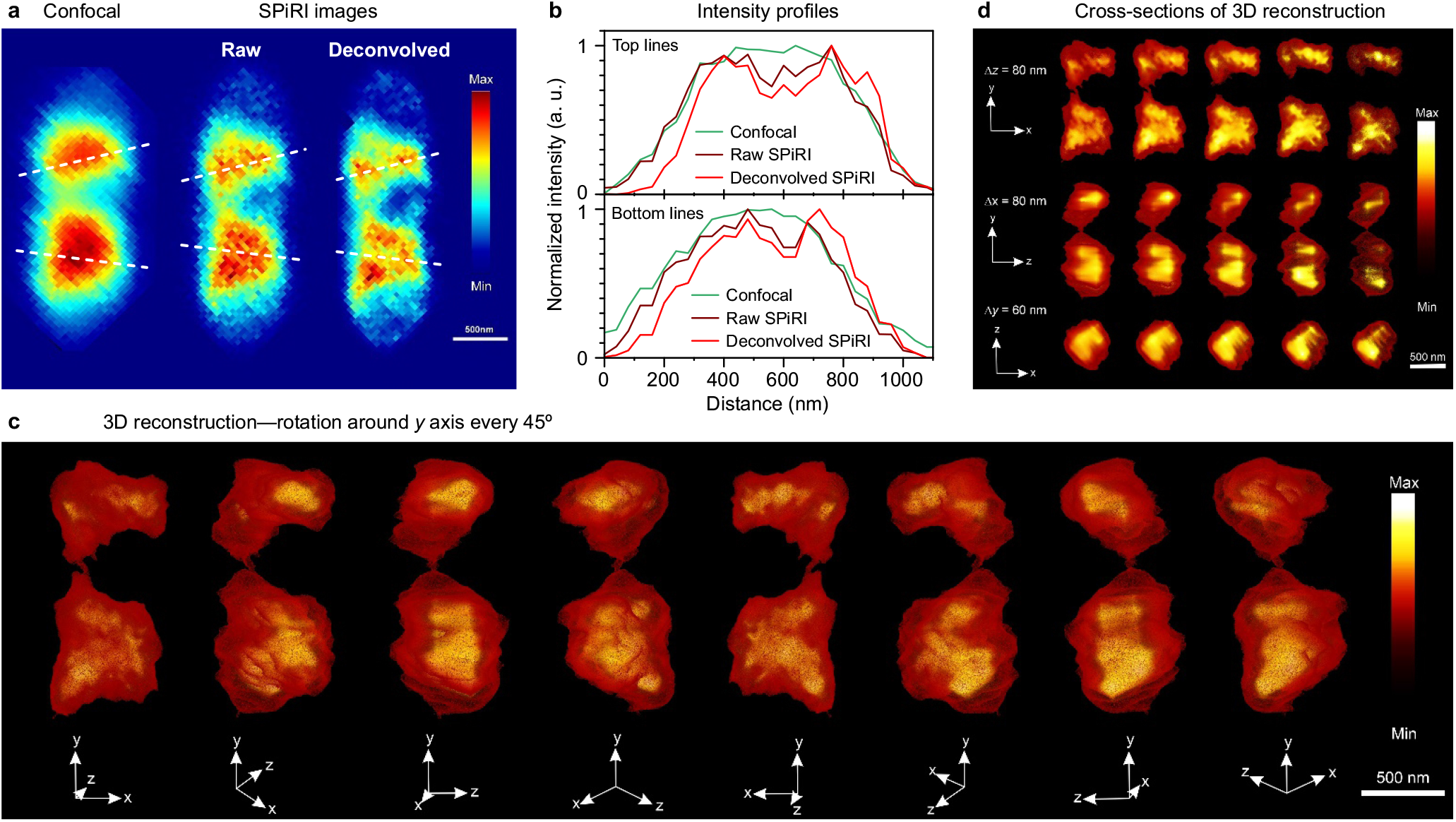
*E. coli* nucleoid stained with SYBR® Gold. **(a)** Reconstructed confocal image and SPiRI images before and after deconvolution. **(b)** Intensity profiles along the white dashed lines in panel (a) for qualitative comparison of the resolution. Intensities of all images were normalized to the 0–1 interval. **(c)** 3D reconstruction from the deconvolved SPiRI images after scanning the whole cell. Each sequential image represents a clockwise rotation around the *y* axis by 45*°* with respect to the previous one. **(d)** Several cross-sections (by 60 or 80 nm steps) from the middle of the 3D reconstruction at *x, y & z* orientations of the cell. The sample was scanned with 40-nm *x & y* steps, and 200-nm *z* steps, *λ*_exc_ = 488 nm, emission collected within 505–545 nm window. 3D reconstruction was obtained from 12 planes.

The outstanding axial resolution of the SPiRI approach allowed us to construct a volumetric 3D image of the *E. coli* nucleoid (see Fig. 5c and Supplementary Movie S1). Several cross-sections of the reconstruction are shown in Fig. 5d (sectioning of the whole 3D reconstruction can be seen in Fig. S7). The resulting nucleoid structure exhibits two longitudinal high-density regions as described previously, but the shape of which appear much more clear in the SPiRI images^36^. Substructures can be observed in each of the high-density regions (Fig. 5c), cylindroids, that constitute subdomains appear clearly. These SPiRI images reveal the 3D structure of the bacterial chromosome with unprecedented precision. The highly structured architectures vary from cell to cell, depending on their growth phase stage. However, common features are maintained, such as elongation of the nucleoid throughout the cell, its cylindrical structure parallel to the cell wall^37–39^, and its global twisting that was also documented before by BALM and PAINT methods^37,38^, although, none of these methods allowed 3D reconstruction of the whole nucleoid in the cell.

## VII. CONCLUDING REMARKS

Several methodologies may nowadays provide theoretically diffraction-unlimited resolutions. Experimental constraints, however, often provide limitations to their resolving power, as well as to their general applicability (special requirement of fluorophores, insufficient photon budget, etc.)^9,40^. Additional restrictions are induced by the use of non-trivial computational methods, which may lead to a risk of artefacts, or to the practical impossibility of generating 3D structures^41,42^. In the present state-of-the-art, a combination of different approaches is still necessary for achieving the most precise description of nanoscale structures^8,43^. Such drawbacks complicate and hinder the application of fluorescence nanoscopy in biological and medical research, which is unfortunate as high-resolution fluorescence imaging has become an essential tool for characterizing nanostructures and their functions.

The method described in this paper, SPiRI, adopts the advantages of ISM, yet avoids sophisticated treatment of the recorded data, such as pixel reassignment (as in ISM), or higher-order statistical analysis (as in SOFI and its variants). In SPiRI, the intensity of only the central pixel of the camera is used for image construction. It results in a significant gain in resolution in all three dimensions, and a very good signal-to-noise ratio. It is worth noting at this stage that once the optical system is well-aligned and considering that *ca*. 3 pixels in each direction correspond to the FWHM of the PSF, the number of photons reaching the central pixel is about 10% of the total photons emitted by the sample. The gain in resolution is thus achieved at the expense of a minimum signal loss.

Implementing SPiRI onto a classical fluorescence microscope, using 488 nm excitation and recording the green region, has allowed us to achieve resolution of 180 and 330 nm in the lateral and axial directions, respectively, without any post-processing of the recorded data. Meticulous calibration of the optical system is predicted to further push the lateral resolution to its theoretical limit of 126 nm under the same excitation conditions. As SPiRI methodology is fundamentally confocal fluorescence microscopy pushed to its limits (actually confocal fluorescence microscopy using a nano-sized detector), the generated images are compatible with the any classical image post-processing techniques, like deconvolution using the experimentally measured PSF of the optical system. Improved resolution in all directions leads to low lateral-to-axial resolution ratio, and results in considerable improvement for reconstructing precise 3D volumes. Other nanoscopy techniques exhibiting an excellent axial resolution exist, however, they are either complex to implement or limited in terms of depth^44,45^, while SPiRI can be implemented in common laser-scanning fluorescence imaging systems. From an experimental point of view, the SPiRI approach is not more demanding than classical confocal microscopy, neither in terms of optics or laser excitation intensity, nor in terms of sample preparation, and can thus be applied to any class of fluorophore and to a wide range of samples of different sizes, fixed or living.

Finally, it must be noted that at higher resolution than 200 nm, optical microscopy characterization at the nanometer range is restricted to fluorescence, as existing super-resolution methods cannot be applied to other emission types. The SPiRI approach being entirely grounded to a quite simple geometry of the signal acquisition and not generating large losses of signal intensity, may be used to generate Raman images at similar resolutions, just by introducing a modification of the emission wavelength selection. A SPiRI Raman microscope is being constructed on this basis, which will open the way for a high-resolution mapping of nanoscale structures containing precise chemical information.

## ACKNOWLEDGMENTS

This work was supported by the European Research Council (ERC) through an Advanced Investigator Grant, contract no. 267333, PHOTPROT (C.I., A.Ga., B.R, D.F.); the platform of Biophysics of I2BC supported by French Infrastructure for Integrated Structural Biology (FRISBI) ANR-10-INBS-05-05 (S.S., C.I., A.Ga., B.R, D.F.); the EU Horizon 2020 research and innovation program under the Marie Skłodowska-Curie, grant agreement no. 675006 (S.S, A.Ga., C.I, B.R.); and the Franco-Lithuanian collaboration Gilibert project (J.C., A.Ge., L.V., S.S., C.I., A.Ga., B.R.) co-funded by the Lithuanian Research Council (no. S-LZ-19-3) and Campus France (no. 42136WA).

## AUTHOR DECLARATIONS

### Competing interests

D.F., A.Ga. and C.I. are shareholders of the Linseg Tech SAS, which is developing a nanoscope based on SPiRI.

### Author contributions

D.F. and A.Ga. conceived idea and built the experimental setup, B.R. and L.V. supervised the project, R.v.G. provided valuable discussions, S.S., C.I. and S.R. prepared the samples, S.S. and C.I. carried out the experiments, J.C. provided theoretical description and performed numerical simulations, S.S. and A.Ge. performed image processing, S.S., J.C. A.Ge., A.Ga. and B.R. wrote the manuscript, S.S. and J.C. prepared figures for the manuscript. All authors discussed the results and commented on the manuscript.

## DATA AVAILABILITY STATEMENT

The data that support the findings of this study are available from the corresponding author upon reasonable request.

## SUPPLEMENTARY MATERIAL

See supplementary material for the description and simulations of the oversampling effect and SPiRI images for variously sized beads, experimentally obtained images of 47-nm sized beads, statistics on bead measurements, and images of *E. coli* nucleoid scanned in *z* direction as well as its volumetric cross-sections (Table S1 and Figures S1–S6).

Movie S1 shows volumetric 3D reconstruction of the *E. coli* bacterial chromatin stained with SYBR® Gold—a rotation around *y* axis. The sample was scanned with 40 nm *x & y* steps, and 200 nm *z* steps, *λ*_exc_ = 488 nm, emission collected at 505–545 nm window. 3D reconstruction was obtained from 12 planes.

## Notes

https://doi.org/10.6084/m9.figshare.21518001.v1

