## Supporting Information for "Single Pixel Reconstruction Imaging: taking confocal imaging to the extreme"

**This PDF file includes:**

Table S1
Figs. S1 to S6
Caption for Movie S1

**Other Supplementary Materials for this manuscript include the following:**

Movie S1

### Image formation

**Oversampling.** Experimentally, the sample is moved across the focused laser beam with a predefined fixed step size, which determines the resolution of the final reconstructed image, *i.e.* the total number of pixels. The numerical simulations of the obtained images of a single and two closely located point emitters for excitation and emission wavelengths being $\lambda_{\mathrm{exc}} = 488\,\mathrm{mm}$ and $\lambda_{\mathrm{em}} = 520\,\mathrm{nm}$, respectively (both intensity profiles are assumed to be expressed as Bessel functions given in Eq. 3 in the main text), with a CCD camera pixel size of 60 nm, and various step sizes used for sample movement are shown in Fig. S1. We see that in all the cases the obtained discrete intensity distributions follow the analytical product function of the excitation and emission profiles (Eq. 17, main text) in the case of a single emitter or the sum of two such functions, centered at their corresponding positions in the case of two point emitters (see red lines in Fig. S1). As expected, the latter continuous distributions might be seen as a limiting case of the registered discrete image when step size $\rightarrow 0$. Although reducing the step size (the so-called oversampling) does produce a smoother image, it does not reveal any additional details, so during the actual measurements one should always choose between image quality (its smoothness) and total data acquisition time. Some image post-processing techniques (e.g. interpolation) may be used to mimic an over-sampled image.

**Table S1 |** FWHM values (in nm) obtained by fits with Gaussian function in lateral and axial planes of several 50-nm-diameter beads, scanned along $x$, $y$, and $z$ directions. Scanning step sizes: $x$&$y$ directions—20 nm, $z$ direction—100 nm. Excitation wavelength $\lambda_{\text{exc}} = 488\,\text{nm}$, emission signal was collected in a 505–545 nm window. No background was removed from any of the images (background removal would result in slightly smaller FWHM values).

| No | # of scanned $z$ planes | SPiRI | | | Confocal | | |
|---|---|---|---|---|---|---|---|
| | | $FWHM_x$ | $FWHM_y$ | $FWHM_z$ | $FWHM_x$ | $FWHM_y$ | $FWHM_z$ |
| 1 | 9 | 165 | 182 | 329 | 216 | 302 | 477 |
| 2 | 10 | 179 | 211 | 269 | 240 | 320 | 445 |
| 3 | 10 | 190 | 213 | 295 | 237 | 338 | 467 |
| 4 | 8 | 192 | 195 | 328 | 225 | 300 | 452 |
| 5 | 9 | 178 | 215 | 329 | 221 | 359 | 423 |
| 6 | 11 | 189 | 221 | 333 | 240 | 324 | 479 |
| 7 | 11 | 186 | 186 | 347 | 244 | 265 | 547 |
| 8 | 9 | 177 | 199 | 381 | 220 | 295 | 430 |
| 9 | 10 | 166 | 184 | 391 | 208 | 247 | 685 |
| Average | 10 | 180 | 200* | 334 | 228 | 306* | 489 |

*$y$ axis is elongated with respect to the $x$ axis due to mechanical drift of $x$&$y$ piezo stage.

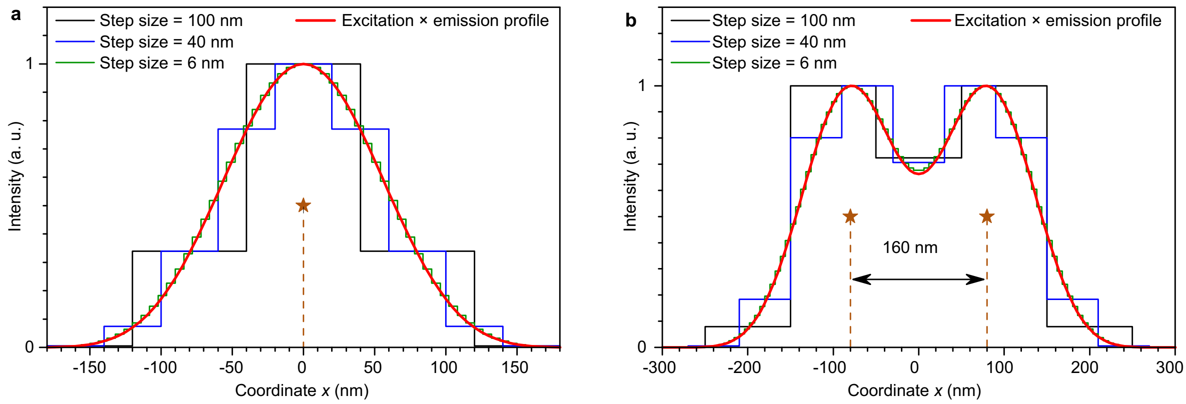


**Figure S1 |** Simulations of oversampling while performing scanning microscopy in the case of (**a**) single emitter and (**b**) two point emitters separated by 160 nm. Simulations were performed using the same parameters and in Fig. 2b and d of the main manuscript ($\lambda_{\text{exc}} = 488\,\text{mm}$, $\lambda_{\text{em}} = 520\,\text{nm}$, NA = 1.45, pixel size = 60 nm, and step sizes are indicated in the legend).

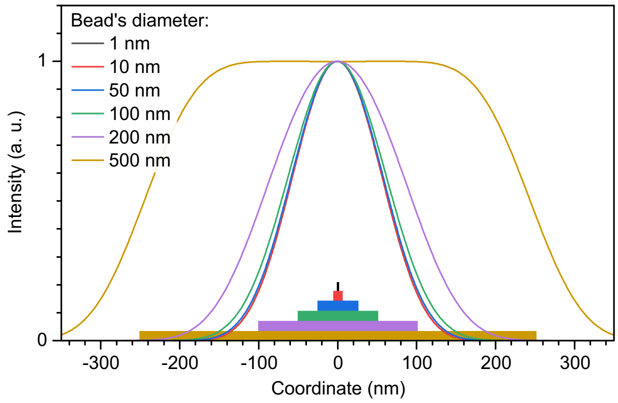


**Figure S2 |** Intensity distribution across the radial cross-section of the simulated SPiRI images that were obtained for the emitting circles of various diameters ("flat beads"). Simulations were performed using the same parameters and in Fig. 2b of the main text ($\lambda_{\text{exc}} = 488\,\text{mm}$, $\lambda_{\text{em}} = 520\,\text{nm}$, NA = 1.45, pixel size = 60 nm). To obtain smooth distributions, very small step size of 1 nm was used. Colored bars at the bottom visualize the size of the corresponding bead.

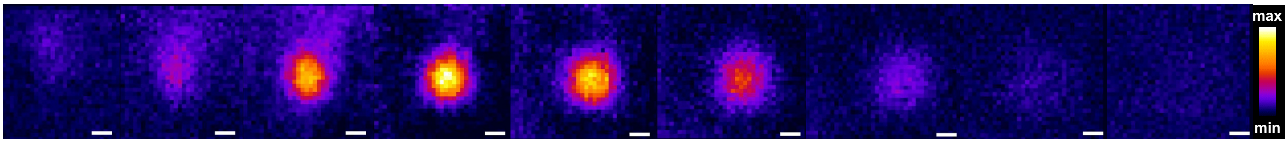


**Figure S3 |** $z$ scans of a 50-nm bead. These images were input for the PSF shown in Fig. 4e of the main text. Scanning step sizes: $x$&$y$ directions—20 nm, $z$ direction—100 nm. Excitation wavelength $\lambda_{\text{exc}} = 488\,\text{nm}$, emission signal was collected in a 505–545 nm window. White bars correspond to 100 nm.

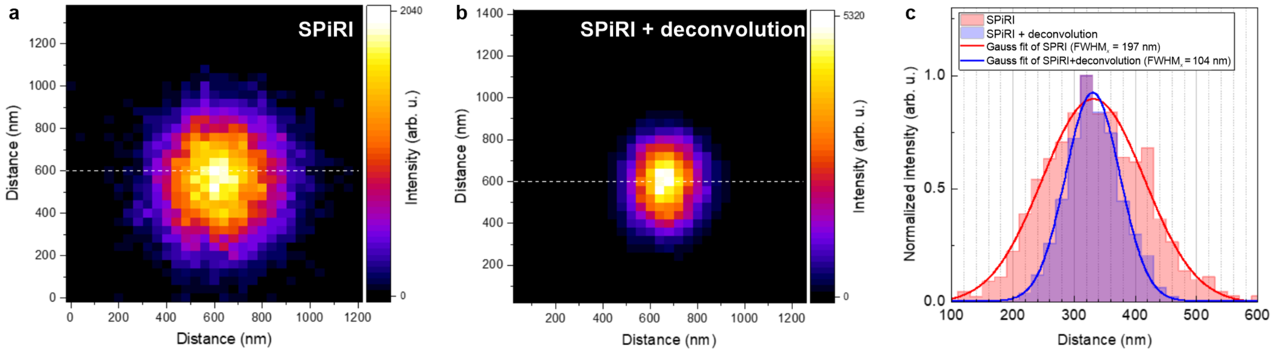


**Figure S4 |** 100-nm bead. **a,** SPiRI image. **b,** Deconvolved SPiRI image. PSF used for deconvolution was obtained experimentally from the 50-nm bead measurements: $\mathrm{FWHM}_{xyz} = 180 : 180\,\mathrm{nm}$. **c,** Comparison of $\mathrm{FWHM}_x$ for SPiRI and deconvolved SPiRI images. The $\mathrm{FWHM}_x$ after deconvolution of SPiRI image of 100-nm bead changes from 197 to 104 nm. Scanning steps for $x$&$y$ directions—20 nm. Excitation wavelength $\lambda_{\mathrm{exc}} = 488\,\mathrm{nm}$, emission collected in a 505–545 nm window.

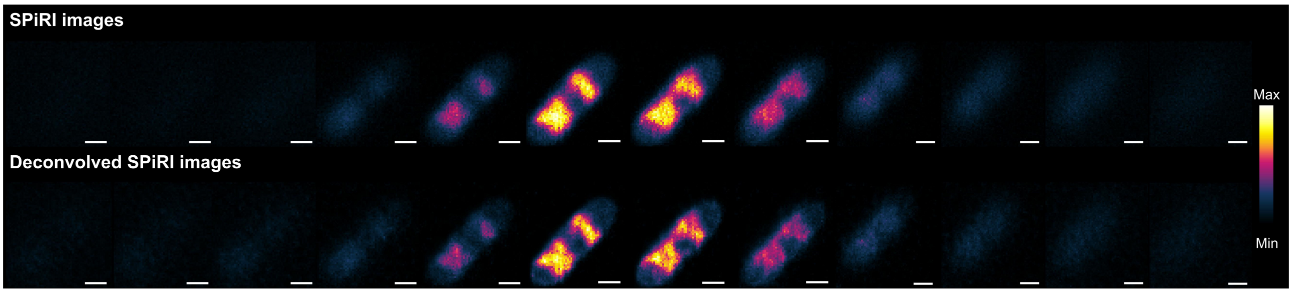


**Figure S5 |** $z$ scans of a *E. coli* nucleoid labelled with SYBR® Gold. **Top,** SPiRI images. **Bottom,** deconvolved SPiRI images. PSF used for deconvolution of each plane was obtained experimentally from the 50-nm bead measurements: $\mathrm{FWHM}_{xyz} = 180 : 180\,\mathrm{nm}$. Scanning step sizes: $x$&$y$ directions—40 nm, $z$ direction—200 nm. Excitation wavelength $\lambda_{\mathrm{exc}} = 488\,\mathrm{nm}$, emission collected in a 505–545 nm window. White bars correspond to 500 nm.

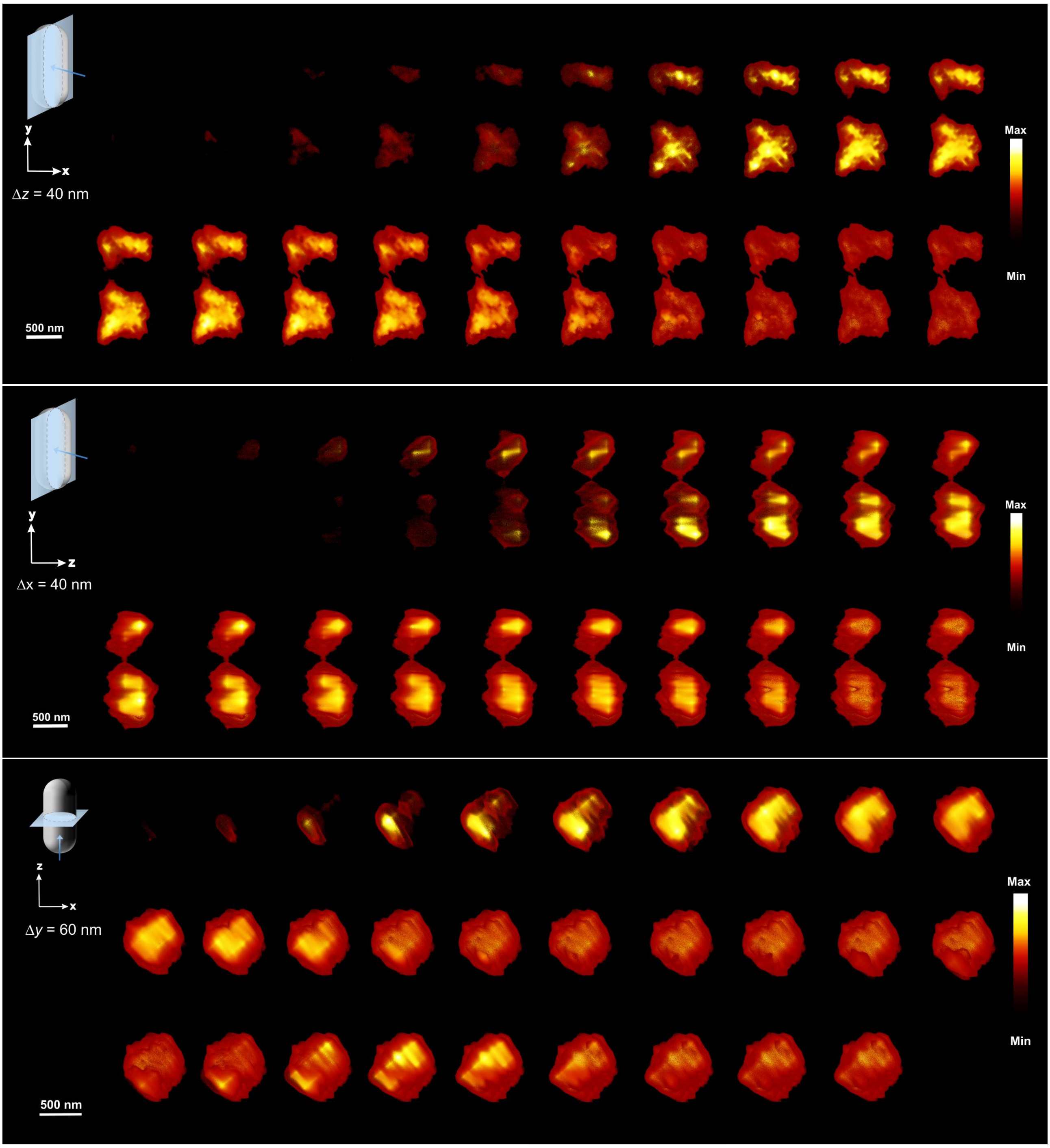


**Figure S6 | ***E. coli* nucleoid labelled with SYBR® Gold. Cross-sections (by 40 or 60 nm steps) of the 3D volumetric reconstruction at $x$, $y$, and $z$ orientations of the cell. Sample was scanned by 40 nm steps in $x$&$y$ directions and 200 nm steps in $z$ direction. Excitation wavelength $\lambda_{exc} = 488\,\mathrm{nm}$, emission collected in a 505–545 nm window. 3D reconstruction obtained from 12 planes.

**Movie S1 |** The movie shows volumetric 3D reconstruction of the *E. coli* bacterial chromatin stained with SYBR® Gold—a rotation around $y$ axis. The sample was scanned with 40 nm $x$&$y$ steps, and 200 nm $z$ steps, $\lambda_{\mathrm{exc}} = 488\,\mathrm{nm}$, emission collected at 505–545 nm window. 3D reconstruction was obtained from 12 planes.
